# Aging-related Transcriptomic Changes with Spatial Resolution in the Human Prefrontal Cortex

**DOI:** 10.64898/2026.01.12.698703

**Authors:** Anupama Rai, Artemis Iatrou, Irais Valenzuela Arzeta, Alexandra L Bartlett, Jane Banahan, Vedant Desai, Lorena Pantano, Isabel Castanho, Pourya Naderi Yeganeh, Athanasios Ploumakis, Shuoshuo Wang, Antonella Arruda de Amaral, Nikolaos Kalavros, Sheethal Umesh Nagalakshmi, Peter Tsvetkov, Sabina Berretta, Isaac H Solomon, Winston Hide, Ioannis S Vlachos, Frank J Slack, Recep Ozdemir, Shannan Ho Sui, Nikolaos P Daskalakis, Maria Mavrikaki

## Abstract

The human prefrontal cortex (PFC), whose laminar organization is essential for cognitive function, is among the first regions to show age-related functional decline^1,2^. Single-cell sequencing studies revealed cell type–dependent aging effects but lacked spatial specificity^3–6^. Spatial transcriptomics (ST) advanced our molecular understanding of the human PFC^7^, yet whether aging-driven changes differ across PFC layers remains unclear. Here, we performed whole-transcriptome ST on postmortem PFC from 37 individuals across the adult lifespan. We mapped cortical layers and revealed aging mechanisms across layers. This represents one of the largest and most comprehensive lifespan ST analysis of the human PFC brain, offering crucial insight into how the brain ages and identifying potential molecular targets to mitigate cognitive aging and extend healthspan.

## Introduction

Extending the adult lifespan is a remarkable achievement of modern medicine, but is accompanied by a rise in age-related conditions, including cognitive aging^8^. This progress paves the way for new frontiers in biomedical research aimed at extending healthy lifespan. Understanding the molecular mechanisms underlying brain aging can guide the development of effective interventions to prevent, delay or even reverse age-related cognitive decline, ultimately extending the productive lifespan.

The human prefrontal cortex (PFC), including the dorsolateral PFC (DLPFC), a region crucial for higher-order cognitive functions and complex behaviors^9^, is particularly vulnerable to age-related changes^1^. It has been reported that aging leads to a reduction of frontal cortex activity^10^, and cognitive decline^11^. On the basis of histological features, the frontal cortex is characterized by six cortical layers (L1-L6) with L2/L3, where most recurrent computations happen, being disproportionally thicker in humans compared to other species^9,12^. Furthermore, these histologically distinct layers also exhibit transcriptional diversity^12^, and neuroimaging studies suggest functional specificity, with upper cortical layers (L1-L3) supporting working memory manipulation and deeper layers (L4-L6) supporting working memory-related responses^13^. Thus, the spatial organization of the human frontal cortex is important for its function and may play a critical role in the aging of different cortical layers.

Advancements in single-cell and spatial technologies have revolutionized our ability to investigate biological processes and diseases at an unprecedented cellular and molecular level^14^. Studies implementing single cell technologies like single cell and single nucleus RNA sequencing have shown that aging induces distinct molecular signatures in neurons and glia in the frontal cortex^6,15^. Although, these studies have provided insights into how different cell types in the brain age, it remains unknown whether their spatial topography in the PFC influences brain aging and function. ST studies in mice indicate that certain brain regions and cells are more affected by aging^16,17^. A recent study explored ST changes in the human PFC of Alzheimer’s Disease (AD) patients and unaffected elderly individuals^15^. Others applied low-resolution ST to examine aging-related changes in the human PFC; however, these studies covered a narrower age range of the adult lifespan and had a limited power to detect spatially distinct transcriptomics changes of aging due to their small sample sizes (up to 22 cases)^18^. In some cases, an inconsistent tissue biopsy was acquired, introducing potential confounds and limiting the consistency of the selected Brodmann area across age groups ^19^. However, a comprehensive spatial transcriptomic study examining the impact of aging on human frontal cortex transcriptome is currently lacking.

In this study, we address this critical gap by leveraging spatial transcriptomics (ST) to investigate the effects of aging on the human DLPFC. We performed an *in situ* transcriptome-wide ST analysis of DLPFC tissues from individuals across the adult lifespan to characterize age-related molecular changes within each cortical layer and white matter (WM). Our findings reveal both shared mechanisms of aging across cortical layers as well as layer specific pathways that contribute to heterogeneity of aging processes. We show aging affects all cortical layers and WM, with L2/L3 exhibiting the most differentially expressed genes (DEGs). Taken together, our study provides the spatially resolved transcriptomic atlas capturing the molecular landscape of aging in the human DLPFC across the adult lifespan, offering unprecedented insights into cortical aging and potential therapeutic targets for cognitive aging.

## Results

### Unsupervised clustering of Spatial Transcriptomics (ST) data reveals laminar-like spatial domains in the DLPFC across the adult lifespan

To employ an unbiased approach in examining how aging affects gene expression profiles in the different cortical layers (L1-L6) of the gray matter (GM) and WM of the DLPFC, we performed whole-transcriptome ST using the Visium v2 platform in DLPFC sections from individuals across the adult lifespan (Fig. 1A, see Table S1A for demographics). Sections with representation of all cortical layers and WM, and adequate anatomical integrity were selected for ST analysis (N=37; see Methods). Each library was sequenced to a median depth of 2.57 x 10^8 reads, with an average of 12,167 Unique Molecular Identifiers (UMIs) and 4,532 genes detected per spot (Supplementary Fig.S1, Table S1B). We filtered out spots with UMIs ≥295 UMIs, resulting in the removal of 6107 spots out of a total of 167613 spots (Table S1C).

**Figure 1:**
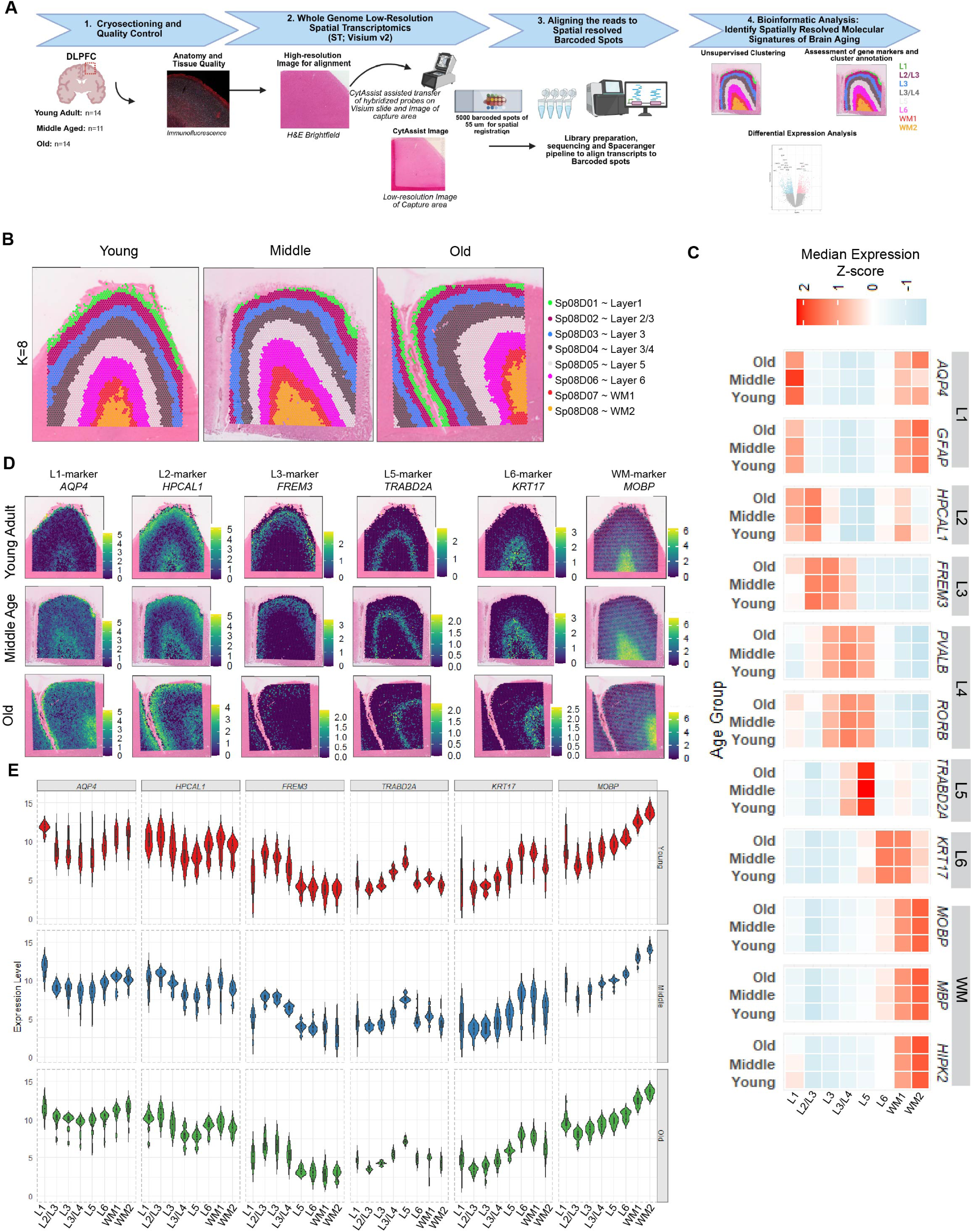
Spatial transcriptomics (ST) study design utilizing the Visium v2 platform to identify aging-regulated genes in distinct spatial domains of the human dorsolateral prefrontal cortex (DLPFC). **A.** Schematic showing an overview of the study design. Fresh frozen DLPFC blocks from individuals across the adult lifespan were included in this study. Cases were grouped into the following three age groups: Young adults, Middle aged, and Old. 10µm thick sections were immunostained with markers that allowed us to ensure the representation of all cortical layers and white matter (WM). GFAP (L1 and WM marker, red), NeuN (neuronal marker and grey matter, white), and DAPI (blue). In a sequential 10µm section which was used for ST, H&E staining was performed and a high-resolution bright field image was obtained. A second low-resolution image of the capture area was obtained by the CytAssist instrument which is used to transfer the hybridized and ligated probes to the barcoded spots on the Visium v2 slides. Probes are extended and eluted to prepare the libraries. Libraries were sequenced and obtained sequences were aligned to the human genome and registered in space using the Space Ranger software (10X Genomics). (Schematic created using Created in https://BioRender.com) **B.** Unsupervised clustering was performed using BayesSpace. Representative samples from the young, middle aged, and old groups showing the laminar spatial organization of clusters at resolution of K=8 (Sp08D01-Sp08D08; top row). **C.** Heatmap illustrating the Z-score of median expression for markers enriched in different cortical layers ^15,22^ in different clusters (Sp08D01:L1-Sp08D08:WM2). **D.** Spot plot showing the expression of previously described cortical layer markers^22^ in the young, middle aged, and old brain. **E.** Violin plot showing the expression of different layer enriched markers in young, middle, and old group presented in D in L1-L6 and WM (1 & 2).

To identify the spatial domains in our samples, we employed the graph-based clustering approach, *BayesSpace*^20^, as it was shown to identify the spatial domains most consistent with the histological cortical layers^7,21^. In our dataset, *K=8* clustering generated more consistent clusters across our samples compared to *K=7* and *K=9*. We also performed unsupervised clustering at K=*16*, but found that some of the clusters lacked clear laminar distribution (Supplementary Fig. S2B); thus, we prioritized *K=8* clustering (Fig. 1B, Supplementary Fig. S2A) for downstream analyses.

To annotate the clusters (Spatial *K=8* Domains 01-08, Sp08D01-08) in relation to classical cortical layer enriched markers (Fig. 2D, 2E) we used established layer-enriched markers obtained from ST study of DLPFC^22^. To further validate our annotations, we closely looked at the expression pattern of layer enriched markers from different studies^5,15^ and we annotated Sp08D01 to L1, Sp08D02 to L2L/3, Sp08D03 to L3, Sp08D05 to L5, and Spo08D06 to L6 (Fig. 1C, Extended Data Fig. 1A, Table S2). Sp08D04 was enriched with L4 markers like *RORB* and L3 markers like *FREM3* and *COL5A2* (Extended Data Fig. 1C, 1D and Table S2), and was thus annotated to L3/L4. We identified two spatial clusters, Sp08D07-08, aligning with WM markers (*MOBP*, *MBP)* (WM1 and WM2; Fig. 1C, Extended Data Fig. 1A, and Table S2). Spatial plots for layer specific markers revealed their spatial distribution pattern across the adult lifespan (Fig. 1D, 1E and Supplementary Fig. S3-S8).

**Figure 2:**
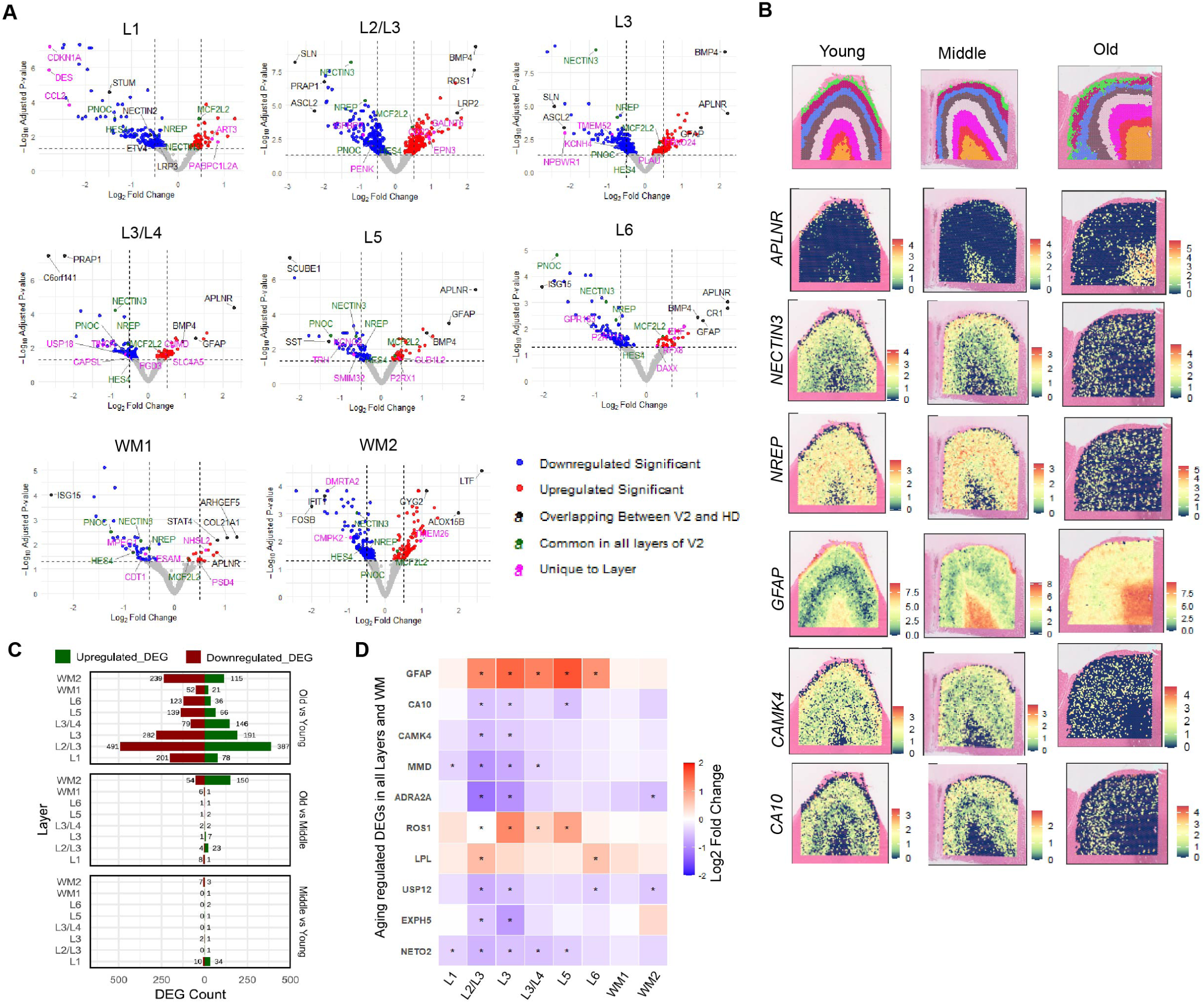
Identification of spatially distinct aging-regulated genes in the human DLPFC. **A.** Volcano plots showing differentially expressed genes (DEGs) in L1-L6 and in two spatial domains corresponding to white matter (WM1 and WM2) based on the old vs. young comparison. Red points, significantly upregulated genes in the older adults (padj < 0.05). Blue points, significantly downregulated genes in the older adults (padj < 0.05). Grey dots represent non-significant genes (padj > 0.05). **B.** Tissue spot plot of genes overlapping between Visium v2 and Visium HD, appearing commonly dysregulated in all the layers (*NECTIN3* and *NREP*), and aging regulated genes reported in earlier studies (*GFAP*, *CAMK4*, *CA10*)^6,23,24^. **C.** Bar graph showing the number of differentially up- and down-regulated genes in different cortical layers (L1-L6) and white matter (WM1 and WM2) in old vs young, old vs middle aged, and middle aged vs young group comparisons. **D.** Heatmap showing the log2 fold-change of aging-regulated DEGs in old vs young individuals across distinct cortical layers and WM, previously identified in bulk RNA-seq and microarray aging studies^6,11,23,24^. * Indicates padj ≤ 0.05.

### Identification of spatially distinct aging-regulated genes in the human DLPFC

To determine whether aging has uniform effects or exhibits cortical layer-dependent and spatially specific influences on the human DLFPC transcriptome, we performed differential expression (DE) analyses across individual cortical layers and WM clusters (Table S3A-S3X), prioritizing comparisons between older adults (>70 years) and younger individuals (<40 years) following an age grouping scheme consistent with our previous studies^23^.

We observed that aging induced transcriptomic changes in all the cortical layers and in the WM (Table S3A-S3H). However, L2/L3 disproportionally thicker in humans ^9,12^ exhibited the largest number of statistically significant differentially expressed genes (DEGs; 878) with padj < 0.05. In contrast, WM1 (Sp08D07) was the least affected by aging, displaying the fewest number of DEGs (73 with padj <0.05) (Fig. 2C, Table S3A-S3H). We further observed that each layer had a unique set of statistically significant aging-related DEGs. L2/L3 had the highest number of unique DEGs (391 genes) followed by L1 with 145 genes. Nine genes, including *NREP (*a previously reported downregulated aging-related gene across all cell types^6^), *NECTIN3, and HES4*, were consistently dysregulated (padj<0.05) across all cortical layers (Fig. 2A and B). We further spatially resolved the expression of certain previously described aging-related DEGs from bulk-tissue, microarray and RNA-seq studies^6,23–25^. Accordingly, *GFAP* was upregulated in L2-L6, and *ROS in* L2-5. In contrast, *CA10* was downregulated in L2-L3 and L5, *CAMK4* in L2-L3. (Fig. 2D and B). We also performed DE analyses comparing old (>70) vs. middle-aged (41–69) and middle-aged vs. young (<40) groups, and found limited transcriptomic changes compared to the old vs. young comparison. (Fig. 1C, Table S3I-S3X).

To characterize the molecular pathways associated with aging across the DLPFC, we performed Gene Set Enrichment Analysis (GSEA) across individual cortical layers (L1–L6) and white matter regions (WM1 & WM2). We identified nine pathways that were commonly dysregulated across all layers and white matter, most notably oxidative phosphorylation (Fig 3 A, B). L2/L3—the region containing the highest number of DEGs—showed dysregulation in pathways such as glial cell differentiation, synapse organization, and the regulation of neurotransmitter secretion (Fig 3C). We also observed significant layer-specific perturbations (padj < 0.05); for instance, L1 exhibited the highest number of unique changes (341 pathways), followed by L3 (64), L2/L3 (60), L6 (20), WM1 (12), WM2 (6), and L5 (5) (Fig 3A).

**Figure 3:**
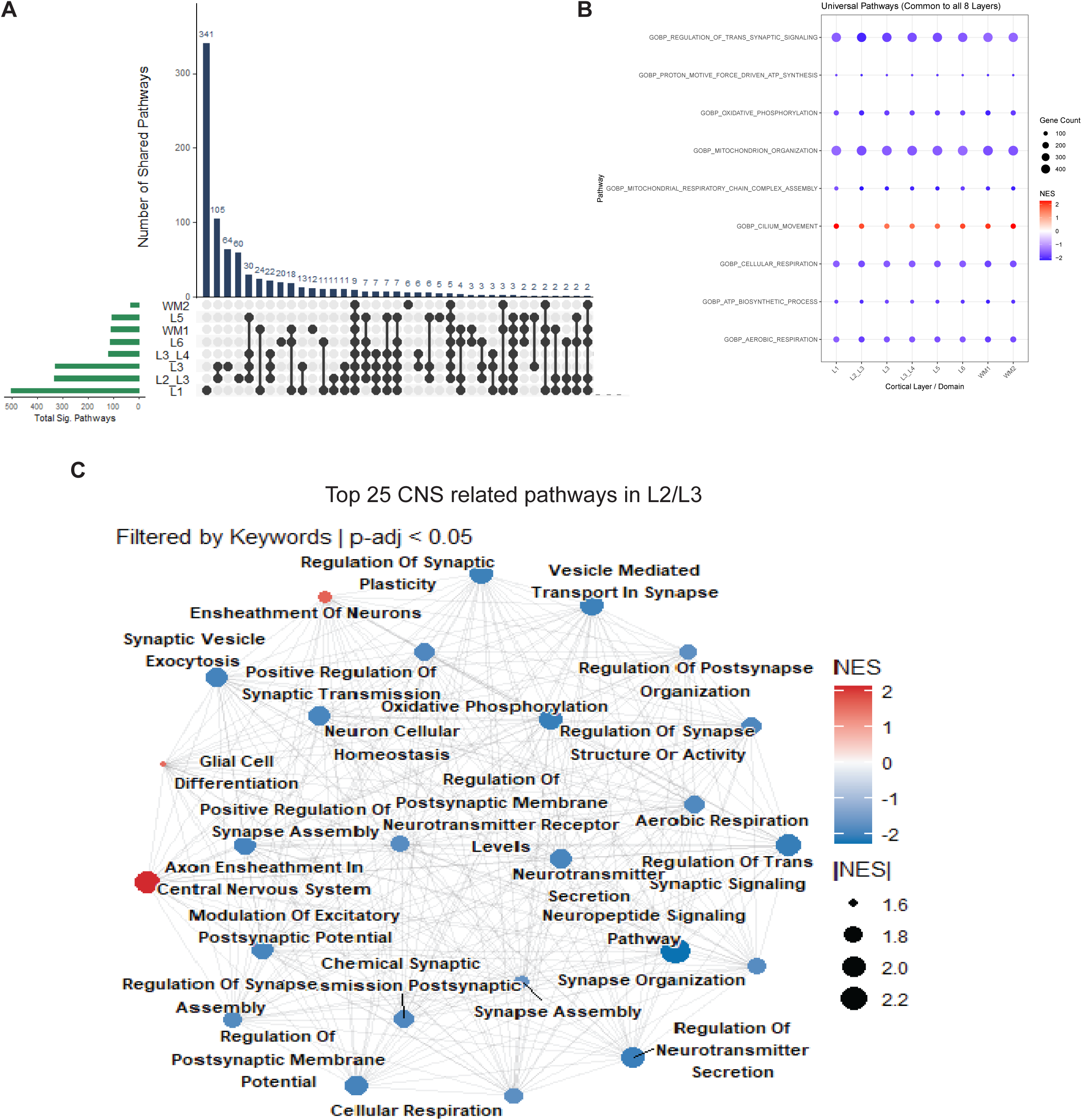
Spatially distinct aging-regulated pathways in the human DLPFC. **A.** UpSet plot showing the overlap in pathways found in old vs young comparison of each cortical layer (L1-L6) and WM. The categories being compared and their size are listed below the bar plots on the left. On the right (directly below each bar) there are dots with connecting lines that denote which categories the overlap is between, or if there is no overlap (just a dot). The numbers at the top of the bars denote the size of the overlap. **B.** GSEA identifies enriched pathways commonly up- and downregulated across cortical layers and WM. Circle size denoted the gene set size and NES score is represented by color scale. **C.** Pathway network map of select top 25 central nervous system (CNS) related pathways in L2/L3.

### Validation of spatially distinct aging-related transcriptomic changes in the human DLPFC using high resolution ST

To further investigate and validate aging-related spatially distinct transcriptomic changes in different cortical layers and WM of the human DLPFC, we adopted the high-definition Visium platform (Visium HD, 10X Genomics), which allows 2 µm resolution, and applied it to a subset of our cohort (N=16, demographics in Table S4A). Data from all 16 libraries (AGHD1-AGHD16) were used to determine the optimal filtering parameters to ensure that we do not discard any biologically relevant data (Fig 4A)

**Figure 4:**
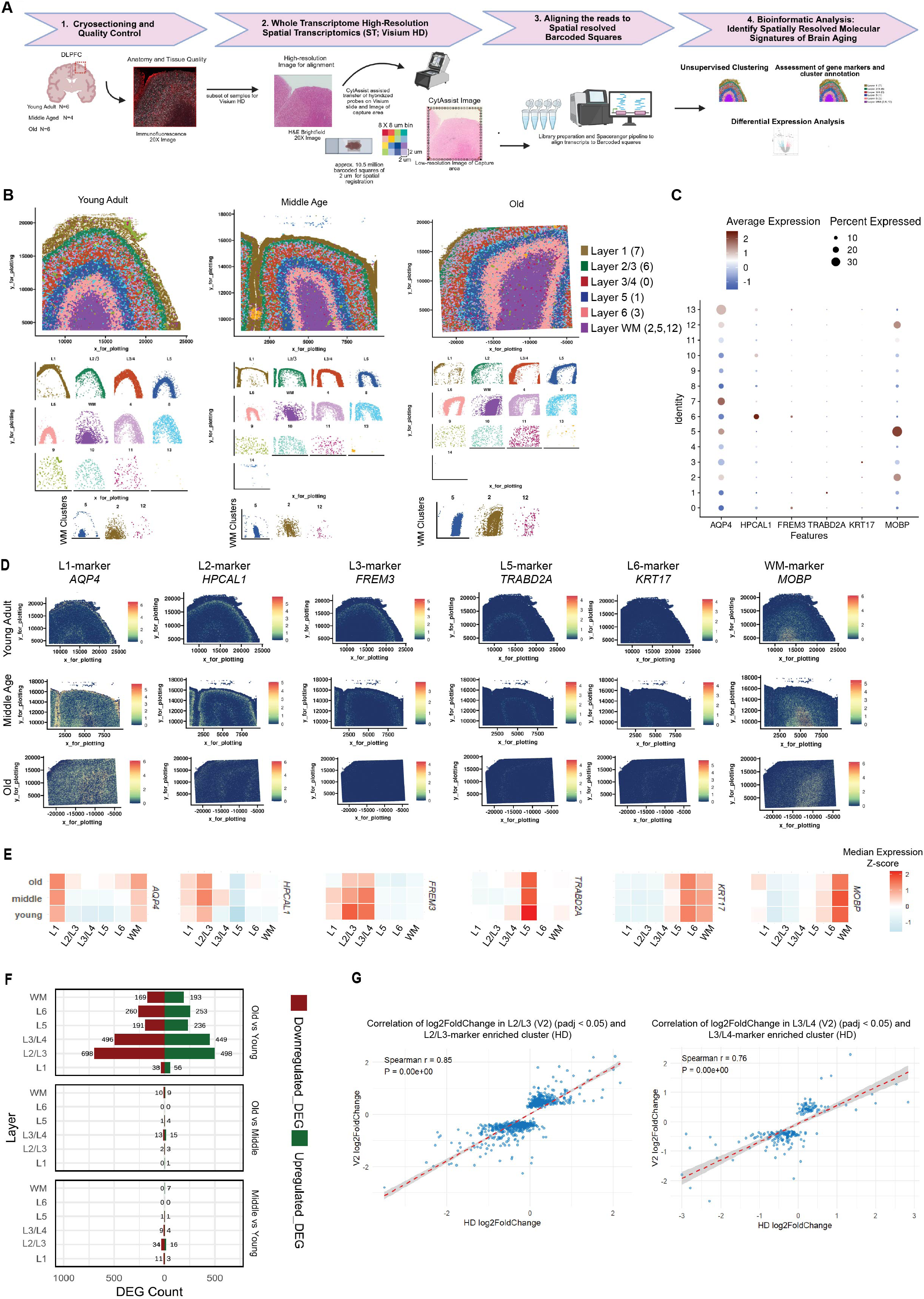
Validation of spatially distinct aging-related transcriptomic changes in the human DLPFC using high-resolution ST (Visium HD). **A.** Overview of high-resolution (2 µm bins). ST approach. A subset of samples (total n=15) from individuals across the adult lifespan (young, n=6; middle aged, n=4; old, n=6) were processed to generate high-resolution ST data using the 10X Genomics Visium HD platform. Immunostaining using GFAP (L1 and WM, red), NeuN (neuronal and grey matter marker, white), and DAPI (blue) was performed to ensure the inclusion of all the cortical layers. In a sequential 10µm section which was used for ST, H&E staining was performed and high-resolution bright field image was taken for each sample to align it with the CytAssist low-resolution image of the capture areas. Hybridized and ligated probes were captured on the HD-visium slides using CytAssist. Probes captured were extended and eluted to prepare libraries. Libraries generated were sequenced and using the 10XGenomics Space Ranger pipeline for HD was used to map the genes to the barcoded bins. Bioinformatic analysis like unsupervised clustering (BANKSY) and differential expression analysis was performed using R. (Schematic created using Created in https://BioRender.com). **B.** Unsupervised clustering using BANKSY in representative young, middle aged, and old samples are shown at the top. X- and Y-shows the co-ordinates. Scatter plot showing the spatial distribution of each individual cluster is shown below. **C.** Dot-plot showing the average expression and percent of bins expressing different cortical layer enriched markers reported earlier ^20^ in BANKSY clusters. **D, E.** Tissue spot plot showing the expression (log count) pattern (**D**) and heatmap of median expression (Z-score) of different layer-enriched genes in young, middle, and old group (Maynard et al. 2021). **F.** Bar chart showing the number of DEGs resulting from pairwise comparison of individual cortical layers in middle aged vs young, old vs middle aged, and old vs young. **G.** Graphs showing Spearman correlation (r) of log2 fold change of aging regulated DEGs (old vs young) for cortical layers (L2/L3 & L3/L4) in Visium v2 (y-axis) and Visium HD (x-axis).

### Unsupervised clustering of high-resolution ST confirms laminar-like spatial domains in the DLPFC across the adult lifespan

To achieve a more spatially informed clustering, we applied the BANKSY workflow^26^, which incorporates not only the expression profile of individual bins but also the mean and gradient of gene expression of their surrounding neighborhoods^26^. Using this approach, we performed clustering in our high-definition dataset and identified clusters which showed more layer-like organization as observed in human DLPFC (Fig. 4B). To annotate the different clusters to cortical layers, we used known layer-enriched markers ^14,22^. We confidently annotated L1 (cluster 7), L2/L3 (cluster 6), L3/L4 (cluster 0), L5 (cluster 1), L6 (cluster 3), and WM (clusters 2, 5, and 12) (Fig. 4C). We further validated our cortical layer annotation using our gene expression profile for each layer generated in the Visium v2 dataset (Extended data Fig. 2B) and using published marker lists^3^ (Fig. 4 D & E). Thus, we concluded that in our high-definition dataset, the BANKSY clustering approach provided laminar-like clusters enriched for cortical layer markers. Tissue spatial plots for different layer-enriched markers revealed their spatial distribution pattern across the adult lifespan (Fig. 4D and Supplementary Fig. S15-S20).

To further investigate aging regulated genes in different cortical layers (L1-L6) and WM of the DLPFC, and validate our Visium v2 findings, we performed DE analysis using BANKSY clusters annotated to cortical layers or WM. We confirmed that there was relatively equal representation of each cortical layer in all the 15 samples, by plotting the number of bins assigned to each layer in each sample (Supplementary Fig. S21). We observed that, with the exception of L1 in sample AGHD2, which contained twice as many bins as other samples due to the inclusion of two L1 regions from adjacent gyri in the capture area, all other layers were similarly represented across samples from different age groups.

Pairwise analysis using DESeq2^28^ confirmed that the largest number of DEGs in each cortical layer occurred between the old vs. young group, as compared to other two comparisons (middle aged vs. young, and old vs. middle aged) (Fig. 4F, Table S5). Furthermore, we observed that similar to our v2 dataset L2/L3 (1196 DEGs) and L3/L4 (945 DEGs) had the most substantial number of DEGs in the old vs young comparison (Fig. 2C and Fig.4F). Interestingly, L2/L3 showed the highest number of DEGs in middle age vs young (50 DEGs), while L3/L4 did so in old vs middle aged (28 DEGs) (Fig. 4F).

### Correlation of low (Visium v2) and high-resolution (Visium HD) ST datasets

We validated our v2 data by performing correlation analysis between the v2 (n=37) and HD (n=15) datasets for our aging-related DEGs for each layer and WM. We observed that the correlation of the two datasets was the highest for L2/L3 (r=0.85) and L3/L4 (r=0.83 for L3, r=0.76 for L3/L4) (Figure 4G) and L5 (r= 0.8). We observed the lowest, yet average, correlation for L1 (r=0.44), which may be partially due to variability of representation of this region in different brain tissues across the larger v2 dataset. For all the other layers, the correlation was strong, approx. r=0.6 (Table S12A-12H). Overall, we concluded that v2 and HD data showed strong concordance in aging-related gene expression changes

## Discussion

Cognitive aging is characterized by extensive transcriptomic changes in the human FC; indeed, and one of the earliest bulk transcriptomic studies of human brain aging revealed substantial gene expression changes emerging after the age of 40^25^, the same threshold defining the young group in our studies. More recent single-cell RNA sequencing studies have shown that these transcriptomic changes of human brain aging are cell type-dependent^4,6^. In parallel, animal studies suggest that aging also induces spatially distinct transcriptomic changes in the mouse brain^26^. However, whether aging-associated transcriptomic changes differ systematically across distinct spatial domains of the human DLPFC remains unresolved.

In this study, we applied ST on postmortem DLPFC sections of individuals across the adult lifespan to assess cortical layer-dependent transcriptomic changes of aging. We provide a comprehensive high-resolution atlas of how the different cortical layers and WM are aging. Our findings reveal shared aging mechanisms across layers, alongside distinct pathways unique to specific layers.

Previous ST studies with a resolution comparable to our Visium v2 studies have shown that unsupervised clustering using Bayespace, a graph-based clustering approach, can reveal laminar spatial domains^7^. Consistent with these findings, applying Bayespace clustering (K=8) in our dataset revealed eight transcriptionally distinct spatial domains across the adult lifespan, including six corresponding to the cortical layers and two corresponding to the WM. By comparing gene expression profiles across these spatial domains, we identified genes altered with aging in all six cortical layers and WM and found that L2/L3 had the highest number of DEGs (old vs. young). Reports indicate that L2/L3 intratelencephalic neurons are expanded in humans compared to other species^9^, suggesting their critical role in complex cognitive functions. Our study highlights the importance of L2/L3 in DLPFC aging. Furthermore, our study also identified substantial number of DEGs in WM, a finding that aligns with established evidence linking age-related cognitive decline to various WM alterations, including increased demyelination, impaired myelin genesis, and axonal degeneration^27^.

As spatial technologies advanced over the course of our study, we utilized high-resolution ST allowing single cell-scale resolution to validate findings from our initial Visium v2 findings in a subset of the cohort. Applying the high-definition ST platform in combination with BANKSY, a spatially aware clustering algorithm^26^, we were able to reveal distinct laminarly distributed spatial domains across the adult lifespan. In agreement with our Visium v2 data, DE analyses confirmed that transcriptomic changes with aging occur across all cortical layers. Pairwise comparisons revealed that the old vs young comparison identified the largest number of DEGs, compared to young vs middle-aged and middle-aged vs old comparisons. This result likely reflects larger effects based on greater differences in age and to some extent greater interindividual transcriptomic variability observed among younger and middle-aged individuals, in contrast to the more uniform transcriptomic profiles evident in individuals over 70 years of age. Notably, gene expression changes were more pronounced in L2/L3 earlier in life, whereas later in life, L3/L4 and L2/L3 exhibited the greatest number of DEGs. Our findings suggest that aging may first affect the upper cortical layers, which predominantly support intracortical connectivity^12^, potentially contributing to subsequent transcriptomic alterations in deeper cortical layers such as L3/L4, L5 and L6. Our analyses identified hundreds of genes exhibiting changes restricted to a single cortical layer, with the largest number detected in L2/L3 (259 DEGs overlapped between HD and v2). Taken together, our study revealed both cortical layer-specific genes associated with aging, and genes likely influenced by layer-independent mechanisms, suggesting the presence of broader molecular programs underlying cortical aging.

## Material and Methods

### Human Postmortem Brain Tissues

Frozen frontal cortex tissue blocks (Brodmann area 8, part of the DLPFC) (received deidentified) from males and females were obtained from the NIH NeuroBioBank (University of Maryland, University of Miami, and Harvard Brain Tissue Resource Center (HBTRC)) and the NIMH Human Brain Collection Core (HBCC) spanning the ages from 18 to 89+ years old. BIDMC IRB approval for permission to conduct research on deceased individuals and receive and report age information for individuals 89+ years old. We have complied with all the relevant ethical regulations. Our criteria for selecting the specimens from the NIH NeuroBioBank were: no clinical brain diagnosis reported, Post Mortem Brain Interval/PMI <48hrs, and reported RNA Integrity Number (RIN) > 7.0. Cases with neurological or psychiatric conditions or COVID-19^23^ were excluded from our study. We also excluded cases with HIV or HEPC due to biosafety reasons. We were collected data from 39 samples which were selected based on the anatomy (inclusion of all cortical layers and white matter/WM), tissue quality (e.g., presence of holes on the sections), and RIN for processing for spatial transcriptomics (ST).

### Cryosectioning

Tissues were shipped from the brain banks on dry ice and were stored at -80°C upon arrival at BIDMC. For sectioning, tissues were kept in the cryostat (Leica CM3050S) at - 20°C for an hour. Chilled OCT (Tissue Tek # MPSMK-981385) was used to mount the tissues on the specimen disc. 10 µm thick sections were obtained using MAX35 or HP35 Ultra microtome blade (Epredia, #3153735) and the Leica cryostat. Sections were stored at -80 °C till further processing.

### RNA Integrity analysis

Tissue punch was collected from specimens, or previously mounted sections were used, to assess the integrity of RNA. RNA was isolated using the Trizol reagent. RIN was assessed using Agilent Tapestation at Biopolymers Facility (BPF Genomics and Core Facility) at Harvard Medical School.

### Immunohistochemistry (IHC)

10 µm thick sections stored at -80°C were used to perform IHC for different markers like CUX2 (upper cortical layer marker), GFAP (marker enriched in L1 and white matter), and NeuN, to ensure the anatomy and presence of L1-L6 and white matter. Briefly, Cryopreserved tissue sections were equilibrated to room temperature (approximately 3 minutes) and subsequently fixed with 4% paraformaldehyde (PFA) for 30 minutes. Sections were then subjected to three 5-minute washes in PBST (1X PBS containing 0.1% Triton X-100, Cat. XYZ). To minimize non-specific antibody binding, sections were blocked with 10% BSA in 1X TBS for 2 hours at room temperature. Primary antibodies, including GFAP (Abcam (ab4674),1:500), NeuN (Abcam (ab177487),1:500), and CUX2 (Thermo Fischer Scientific (BS-11832R-has been discontinued), 1:150), were applied overnight at 4°C in a humidified chamber. The following day, sections were washed three times for 5 minutes each in PBST and incubated with Alexa Fluor secondary antibodies (1:500) (Goat Anti-Rabbit Alexa Fluor 750 (A-21039; Thermo Fisher Scientific), Goat Anti-Mouse Alexa Fluor 594 (A-11005, Thermo Fisher Scientific), and Goat Anti-Chicken Alexa Fluor 647 (A-21449, Thermo Fisher Scientific)) for 2 hours. Sections were counterstained with DAPI (1:1000, Thermo Scientific #62248) and imaged using epifluorescence microscope at 20X magnification on an Akoya PhenoImager HT at BIDMC Spatial Technologies Unit (STU). Based on the anatomy revealed by the IHC images, a 6.5mm x 6.5mm region of interest (ROI) that included all six cortical layers and WM was selected to perform the ST analysis.

### 10X Genomics Visium v2

Was performed in 35 samples at the BIDMC STU. We also run 4 additional samples in-house. All procedures were performed according to manufacturer’s recommendations. Briefly, slides were retrieved from -80°C and equilibrated to room temperature for 5 minutes. Slides were submerged in pre-chilled methanol and incubated for 30 minutes at - 20°C. Slides were stained using Hematoxylin (Millipore Sigma, MHS16) and alcoholic eosin (Millipore Sigma, HT110116; together called H & E staining) were used to counterstain the tissue according to manufacturer’s (10X Genomics) Document CG000614 Rev A. The slides were coverslipped using 80% glycerol solution and immediately scanner for bright-field image, at 20X magnification on an Akoya PhenoImager HT. Coverslips were removed after imaging. Slides were placed in Visium cassettes (10X Genomics, PN-1000519) and treated with 0.1N Hydrochloric Acid at 42°C for 15 minutes to destain the hematoxylin. Samples were immediately taken to probe hybridization.

Following Document CG000495 Rev F (10X Genomics), sections were hybridized for 16 hours at 50°C, using Visium Human Transcriptome Probe Set V2 (10X Genomics, PN-1000466) on a BioRad C1000 thermal cycler with 96-Deep Well Reaction Module for 18-24hrs. After probe hybridization, the slides were washed on the thermal cycler at 50°C, and a ligation reaction was performed. Two washes were performed at 57°C and slides were re-stained with 10% alcoholic eosin solution before loading into the CytAssist instrument (10X Genomics). Tissue removal and probe transfer to the Visium slides were performed using reagents from the FFPE Reagent Kit v2 (10X Genomics, PN-1000436) using CytAssist.

After probe extension and elution, a pre-amplification PCR reaction of 10 cycles was performed and the yielded DNA was quantified using qPCR on a QuantStudio thermal cycler (Applied Biosystems). The readout was used to determine the cycle number for indexing PCR, following the protocol recommendation. Pre-amplified libraries were indexed with Dual Index Plate TS Set A (10X Genomics, PN-3000511). After SPRI beads cleanup, library concentrations were measured on a Qubit Flex with Qubit dsDNA HS Assay Kit, Agilent Fragment Analyzer 5200 (and D1000 ScreenTape), and finally sequenced on a Singular Genomics G4 sequencer (Singular Genomics, San Diego and NovaSeq XPlus) with paired end reads for no less than 25,000 read pairs/spot according to the manufacturer’s guidelines.

### 10X Genomics High-Definition Visium (Visium HD)

We performed high-definition ST on 16 samples, spanning across the adult lifespan. The 16 samples used were a subset of the cohort analyzed in our Visium v2 studies described above. Selection of this subset was based on tissue quality, age distribution, and post-sectioning interval. The protocol provided by 10xGenomics was followed (CG000763 (RevA) and CG000685 (RevB)). Briefly, sections stored at -80°C in slide mailers were thawed in the thermocycler for 1 minute at 37 °C. The sections were fixed in 3.7% Formaldehyde (Thermo Fisher Scientific, BP531-500) for 30 minutes and Hematoxylin (Millipore Sigma, MHS16) & Eosin (Millipore Sigma, HT110116; H & E) staining was performed. We used the lower concentrated Mayer’s hematoxylin which has been standardized for v2 assays. Bright field images were captured at 10x and 20x using Keyence microscope (BZ800 series). Post imaging destaining was performed (0.1 N HCL, Fisher Chemical, SA54-1) and permeabilization was performed using 1%SDS (Millipore Sigma, #71736) and 70% methanol (Millipore Sigma, #34860). Probes for the whole transcriptome (Kit:1000466) was hybridized for 16-18 hours in the thermocycler (BioRad (PTC Tempo Deepwell)) at 50°C. Probe ligation was performed as recommended and CytAssist enabled probe release was done and captured on washed and dried HD visium slides (Kit: 1000669). Captured probes on HD visium slides were extended in the thermocycler and eluted in 0.08M KOH (Millipore Sigma, P4494) and 3ul of 1M Tris-HCL (Thermo Fisher Scientific, AM9855G) was added to each tube. Pre-amplification of the eluted probes was performed in the thermocycler and the product clean-up was done using SPRIselect. To determine the number of cycles for library preparation, real time PCR was performed using the KAPA universal SYBR green (KAPA Biosystems, KK4824) using 1ul of 2:8 diluted pre-amplification product. Following the 10xGenomics recommendations and qPCR amplification plots, cycle number (Cq) was determined. Libraries were prepared following 10X genomics protocol and SPRIselect (Beckman Coulter, B23318) cleanup was performed. Libraries were sent to Azenta for quality control (Qubit and DNA tapestation (D1000 ScreenTape)), quantification (using qPCR) and sequencing. Pooled libraries were sequenced using NovaSeq XPlus.

## Bioinformatic Analysis

### Visium V2 raw data processing

To map the sequencing reads spatial coordinates, we used the *SpaceRanger* software (v2.1.1, from 10x Genomics. Briefly, raw sequencing data files (FASTAQ) were processed with the count pipeline, which aligned the reads to the human reference genome (GRCh38-2020-A) and generated spot-level gene expression matrices. Tissue detection and image alignment were performed using the brightfield histology images and the corresponding spatial barcodes from the Visium platform. The outputs from this pipeline were used for downstream quality control and analysis in R version 4.3.3.

### Quality Control (QC) Parameters for the Visium V2 Dataset

We imported the raw Visium outputs into R using the read10xVisiumWrapper() function from the spatialLIBD package^22,28^, and stored the SpaceRanger output in a SpatialExperiment object containing raw UMI counts, spatial coordinates, and both low- and high-resolution histology images^29^ for all 39 samples. To ensure spatial relevance, we excluded spots located outside the tissue capture area using information from the tissue_positions.csv files^29^, retaining only those within the annotated tissue boundaries.

Spot-level QC was performed using the scran^30^ and scater^31^ packages. We applied the quickPerCellQC() function to compute automatic thresholds for UMI counts, number of expressed genes, and mitochondrial gene content. The derived thresholds were: ≥544 genes per spot, ≥295 UMIs, and ≤15.3% mitochondrial gene expression. Spots with low library size were flagged and removed using the isOutlier() function^31^ on log10-transformed total UMI counts in a sample-aware manner, resulting in the exclusion of 1,889 low-quality spots.

Although the pipeline identified spots with high mitochondrial content, we chose not to exclude them. Prior work has demonstrated that this threshold is not critical in human DLPFC samples^22^ and considering that mitochondrial function is known to decline with aging^32^, we aimed to preserve potentially informative biological signal. A summary of the number of spots before and after filtering is provided in Supplementary Table S1C. The resulting filtered object served as the foundation for all downstream analyses.

We then applied sample-aware normalization using the scran package^30^. We computed size factors with computeSumFactors() after clustering spots within each sample using quickCluster(). The log-transformed normalized expression matrix was obtained via logNormCounts(). To identify highly variable genes (HVGs), we modeled gene-wise variance using modelGeneVar() with sample ID as a blocking factor. The top 10% most variable genes were selected for dimensionality reduction, along with subsets defined by FDR thresholds of 0.05 and 0.01 for exploratory analyses.

Dimensionality reduction was performed using principal component analysis (PCA) on the HVGs. We selected the number of PCs based on an elbow point identified using the findElbowPoint() function from the PCAtools package^31^. To address potential batch effects, we applied Harmony batch correction using sample ID as the integration variable.

UMAP and t-SNE projections were first computed using the uncorrected PCA embeddings (n = 50) via runUMAP() and runTSNE(), with t-SNE run at a perplexity of 80. Subsequently, UMAP embeddings were recomputed based on Harmony-corrected PCs and used in downstream analyses.

### Unsupervised Clustering using BayesSpace of V2 data set

We next used BayesSpace^33^ (for unsupervised clustering of the spots into spatial clusters. We iterated through incremental clustering resolutions (k = 2-16), and selected k = 2,8, and 16 for downstream analysis.

During evaluation of cluster maps, two samples were excluded from the working dataset due to poor spatial structure. One of these showed very low gene detection (860 genes per spot on average), likely due to technical artifacts during library preparation. The second was derived from a homicide case, and presented a disorganized cluster pattern, possibly reflecting underlying biological or environmental disturbances. In both cases, we could not confidently delineate cortical layers or white matter, so these samples were omitted from analyses requiring accurate laminar annotation.

### Visium V2 cluster annotation to frontal cortex cortical layers

To interpret the biological relevance of the spatial clusters, we performed spatial registration to annotated cortical layers using layer-enriched gene expression profiles available in the sce_layer data previously^22^. We used the spatialLIBD package^28^ to implement this analysis.

We focused on the clustering result obtained at resolution k = 8 and loaded the spatial registration enrichment statistics generated for this resolution. From the registration output, we extracted moderated t-statistics (t-stat) for each gene and cluster combination. These t-statistics quantify the enrichment of each gene in each BayesSpace cluster.

To compare our data with publicly available laminar gene expression profiles, we retrieved the reference modeling results using the fetch_data function, which contains gene-level statistics for each of the six cortical layers from prior bulk RNA-seq data of the human DLPFC^22^.

We computed pairwise correlations between the top 100 t-statistics for each BayesSpace cluster and each cortical layer using the layer_stat_cor() function. This approach enabled us to infer the most likely layer identity of each spatial cluster based on transcriptional similarity. The results were visualized as correlation heatmaps.

### Differential Expression (DE) analysis to identify aging regulated genes in V2 dataset

To determine whether different cortical layers and spatial domains exhibit common or region-dependent transcriptomic changes with aging, we performed DE on distinct spatial domains. Specifically, we assessed three group comparisons: old vs. young, middle-aged vs. young and old vs. middle-aged. Pseudobulk aggregation was performed using the registration_pseudobulk() function from the spatialLIBD package, by summing counts across spots within each individual and domain from BayesSpace-defined domains (k = 8) aligning the spatial domains with the corresponding cortical layer^22,28^. The resulting object included the variable age_layer, which encodes the combination of age group (Young, Middle, Old) and BayesSpace domain (Sp08D01–Sp08D08). DE was performed using DESeq2^34^ on pseudobulked count matrices derived from^34^. A DESeq2 dataset was constructed using the design formula ∼ age_layer + sex, and genes with total counts ≤10 across all samples were filtered out^34^. For each of the eight domains, we tested differential expression across three pairwise comparisons: Young vs Middle-aged, Old vs Middle, and Old vs Young. Log2 fold changes were moderated using the adaptive shrinkage estimator (type = “ashr”), and results were annotated with gene symbols. Differentially expressed genes were exported by domain and comparison for downstream analysis.

### Visium HD raw data processing

Visium HD raw sequencing files (FASTQ) for 16 samples were processed using the 10xGenomics Space Ranger online platform on the 10X Genomics icloud. Automatic alignment (or manual alignment when needed) of high-resolution H&E bright-field image with CytAssist captured image was performed using Loupe browser (v8.1.2; 10xGenomics).

### Unsupervised Clustering using BANKSY to identify spatially resolved clusters in HD data set

We performed clustering using BANKSY^26^, which takes into account not only an individual bin’s expression pattern but also the mean and the gradient of gene expression levels in a bin’s broader neighborhood. This makes it valuable for identifying and defining spatial regions of interest. We used parameters lamba = 0.8 and k_geom = 8. Sample AGHD16 did not show clustering consistent with other samples, and thus was excluded from DE analysis.

### Pathway enrichment analysis in aging

To identify cellular processes and biological pathways associated with aging in our Visium v2 we used Gene Set Enrichment Analysis (GSEA) for each cortical layer and cell type in old vs young, ranking DEGs by log fold change.

## Data and code availability

Visium v2 and HD data will be deposited on GEO and will be made publicly available upon publication. Codes used will be available on GitHub repository upon publication.

## Author contributions

MM conceptualized and designed the study, acquired funding, and supervised the project. AR and MM coordinated the project. AR prepared and selected tissues, conducted a subset of Visium v2 experiments. SW, AP, and AADA performed the Visium v2 experiments under ISV’s supervision. SUN and NK generated Spaceranger files for the Visium v2 dataset, supervised by ISV. AI and IVA carried out bioinformatics analyses of the Visium v2 dataset, supervised by NPD, AI and MM. AB under SHS’s supervision performed pilot bioinformatic analysis. PNY, IC, LP, PT, WH, and RO advised on the analysis. IVA, AI, AB, AR, and VD generated the plots. FS contributed to funding acquisition and provided advisory support. SB provided suggestions for tissue selection. IHS contributed to funding acquisition and provided tissues for pilot studies. JB assisted with tissue preparation and quality control assessment. AR and MM wrote the manuscript. All authors reviewed, edited, and approved the final manuscript.

## Disclosures

NPD has been on scientific advisory boards for BioVie Inc., Circular Genomics, Inc., and Polaris Genomics, Inc. for unrelated work. MM consults for ExQor Technologies and ISV consults for AlteRNA Therapeutics, Chronicle Medical Software, Guidepoint Global, and Mosaic for unrelated work. All other authors declare no conflicts of interest.

## Acknowledgements

This study was supported by an R01 grant (5R01AG079799 to MM) from the National Institute on Aging (NIA). We thank the NIH NeuroBioBank (University of Maryland, University of Miami, and Harvard Brain Tissue Resource Center; https://neurobiobank.nih.gov/) and the Human Brain Collection Core of the NIMH Intramural Research Program (http://www.nimh.nih.gov/hbcc) for providing the tissues analyzed in this study. We also thank Stefano Marenco, Alexandra LeFerve, and David Davis for assistance with the identification of appropriate tissues, and the Harvard Biopolymers Core for performing quality control on our samples. We thank Stephen Wood from 10x Genomics for advising us on the Visium pipelines. AI was supported by 2023 Young Investigator grants from the Brain & Behavior Research Foundation to AI. NPD was supported by 2015 and 2018 NARSAD Young Investigator grants from the Brain & Behavior Research Foundation to NPD. AI and NPD were supported by a R01MH133268 from the National Institute of Mental Health to NPD. We thank the Dana-Farber/Harvard Cancer Center in Boston, MA, for the use of the Spatial Technologies Unit (RRID:SCR_024905), which provided 10x Genomics V2 Analyses. Dana-Farber/Harvard Cancer Center is supported in part by a NCI Cancer Center Support Grant # NIH 5 P30 CA06516. FJS was supported by 5R01AG079799 and R01AG082093-01 and WH was supported by R01AG082093-01. The authors acknowledge the use of the BIDMC Ithaca High Performance Computing Cluster of the Spatial Technologies Unit, which supported a portion of the analyses. We also acknowledge the use of the Harvard Medical School High Performance Computing Cluster (O2).

